# Extreme temperatures impede the release success of captive-bred avian scavengers

**DOI:** 10.1101/2024.03.14.585025

**Authors:** Nili Anglister, Marta Acácio, Gideon Vaadia, Eitam Arnon, Michael Bruer, Ohad Hatzofe, Ygal Miller, Roni King, Noa Pinter-Wollman, Orr Spiegel

## Abstract

Conservation translocations (reinforcements and reintroductions) are central for managing various endangered species, yet, their implementation is logistically and financially challenging. Because many translocations fail due to the mortality of released individuals, identifying and preventing these factors is crucial. Here we examine risk factors affecting the post-release survival of the Griffon vulture (*Gyps fulvus*). The Israeli population is facing extinction, and the recovery efforts by the local Nature and Parks Authority include supplemental releases of individuals from captive-breeding or rehabilitated birds (mostly imported from Spain). We use GPS tracking and thermometers to compare movement, behavior, and weather conditions experienced by released and wild-born individuals active in the same area and time. Our results show that the Judean Desert release site had a significantly lower survival rate than other release sites included in the program. After excluding several possible factors (e.g. known pathogens) with pathological examinations, we found that released individuals at this area were exposed to higher temperatures preceding their death (compared to wild-born griffons nearby), suggesting heat stress as the cause of death. Arguably, they failed to avoid the harsh environmental conditions, resulting in overheating due to their inexperience and undeveloped flight ability, as reflected by their lower probabilities of flying and shorter travel distances per day compared to wild-born vultures. These findings have led to adjustments of the local release protocol, (namely winter releases) resulting in a significant improvement in the early survival of translocated Griffons in the Judean Desert. Accounting for the harmful effects of extreme weather conditions is particularly important in a world facing climate change. More broadly, this sequence of scientific investigation with data integration from ecological, clinical, and biotelemetry sources, leading to improvement in translocation success demonstrates how conservation practices can be optimized by supporting studies to ensure the survival of endangered species in the wild.

## 1. INTRODUCTION

Conservation organizations have tried to stop rapid species declines for over a century. One of the most popular methods used to recover locally endangered or extinct species is the conservation translocation of animals to reintroduce to their historic range, or reinforce endangered populations with individuals from other locations or breeding programs (Griffith et al., 1989; Ripple & Beschta 2012; IUCN/SSC 2013; Corlett 2016). For simplicity, hereafter we use ***translocation*** as a general term for these diverse conservation methods. These projects require massive efforts and financial investment in captive breeding, release, and monitoring of the translocated individuals, yet many of these projects have been unsuccessful (Wolf et al., 1996; Fischer & Lindenmayer 2000; Bubac et al., 2019; Morris et al., 2021). Unfortunately, the outcomes of translocations are not always well documented and shared, especially when they are unsuccessful (Scargle 2000; Schooler 2011; Berger-Tal et al., 2020), hindering evidence-based conservation (Fischer & Lindenmayer 2000; Sutherland et al., 2004; Seddon et al., 2007).

The survival of released individuals is a function of the animal’s internal condition and their interaction with the external, biotic and abiotic conditions (Maran et al., 2009; Hardouin et al., 2014; Tarszisz et al., 2014; Bellis et al., 2020). For example, the survival of translocated individuals can be impacted by the physiology and morphology of captive-born individuals, as well as their early life experiences (or lack of), compared to native wild ones (Jule et al., 2008; Tarszisz et al., 2014). Captive-bred individuals may also be immunologically naïve to local pathogens or lack knowledge about local threats and predators, which might have caused the species’ decline (Williams et al., 1988; Stanley-Price 1989; Bubac et al., 2019; Greggor et al., 2019). Recently translocated individuals also lack information about resources, such as food and shelter, necessary for their survival in the new environment (Pinter-Wollman et al., 2009; Maor-Cohen et al., 2021). Finally, translocated animals may be exposed to environmental conditions they are not accustomed to, which can be harsh for wild-born animals (Altwegg et al., 2006; Nevoux et al., 2008; Strandberg et al., 2010) and even more for captive-bred, reintroduced individuals (Nicoll et al., 2003; Tavecchia et al., 2009; Hardouin et al., 2014).

Consequently, mortality is often very high post-release, especially during the first month in the new environment (Calvete & Estrada 2004; Maran et al., 2009; Tavecchia et al., 2009; Devineau et al., 2010). Post-release movements can determine the survival of translocated animals by shaping how they acquire information or encounter risks. Exploratory movements are very common in many taxa (Gouar *et al.,* 2008a; Linklater & Swaisgood 2008; Pinter-Wollman 2009; Marques et al., 2011; Nussear et al., 2012; Pille et al., 2018; Griffith et al., 2019; Bilby & Moseby 2023). However, some animals remain in the vicinity of the introduction area post-translocation (Nafus et al., 2017; Garnier et al., 2021). Compared to wild-born animals, captive-bred animals (raised in constrained conditions, such as aviaries) tend to remain close to the release site due to reduced physiological abilities (e.g. poor flight capacity) (Putaala et al., 1997; Hess et al., 2005; Whiteside et al., 2016; Lopes et al., 2018; Franzone et al., 2022). Given the high investment in released animals, and their enhanced risk, an adaptive management approach, with real-time monitoring of released animals, can greatly increase translocation success. Tracking released animals with transmitters (GPS or radio telemetry) offers important data on survival, reproduction, habitat use, and movement patterns. These benefits are especially pronounced for highly mobile species covering large areas that make direct observations less practical (Maran et al., 2009; Pinter-Wollman 2009; Devineau et al., 2010; Efrat et al.,, 2022).

Vultures are large obligate scavengers covering vast areas in search of carcasses, making them particularly sensitive to human-related threats. Indeed, vulture populations are collapsing worldwide with 17 of their 22 species in decline (Ogada et al., 2012; Buechley & Şekercioğlu 2016). Nevertheless, active conservation can counteract this trend, and effectively increase population size demonstrating the importance of this approach. For instance, conservation of Griffon vulture (*Gyps fulvus* hereafter griffons) has resulted in positive outcomes in Western Europe (Sarrazin et al., 1994; Gouar et al., 2008a; Birdlife International 2020). In Israel, in contrast, the situation is less positive; three out of five vulture species that had local populations historically are already locally extinct as breeders, and the remaining two - Egyptian (*Neophron percnopterus*) and griffons are critically endangered (Mayrose et al., 2017; Birdlife International 2020). The griffon population in Israel declined rapidly - from thousands of individuals a century ago, to approximately 500 individuals a couple of decades ago, to less than 200 individuals today. This decline is caused by poisonings, collision with infrastructure and electrocutions, habitat loss and low reproductive success rates (Mayrose et al., 2017; Choresh et al., 2019; Anglister et al., 2023). To address this decline, the Israeli Nature and Parks Authority (hereafter, INPA) runs an extensive management program, that includes: A. Operation of supplementary feeding stations & sanitation of livestock carcasses; B. Population monitoring (annual surveys, morbidity and mortality analyses, and tagging griffons with wing tags, rings, and GPS transmitters); C. Conservation translocations of vultures born in Israeli captive breeding programs, as well as vultures imported from Spain to three main sites: the Carmel mountain, the Golan heights (both in the north of Israel with a Mediterranean climate) and the Judean desert in the south of Israel (with a desert climate) (**Fig. 1**). Here, we focus on examining the success of the conservation translocation of captive-bred and imported griffons in Israel.

**Fig. 1.**
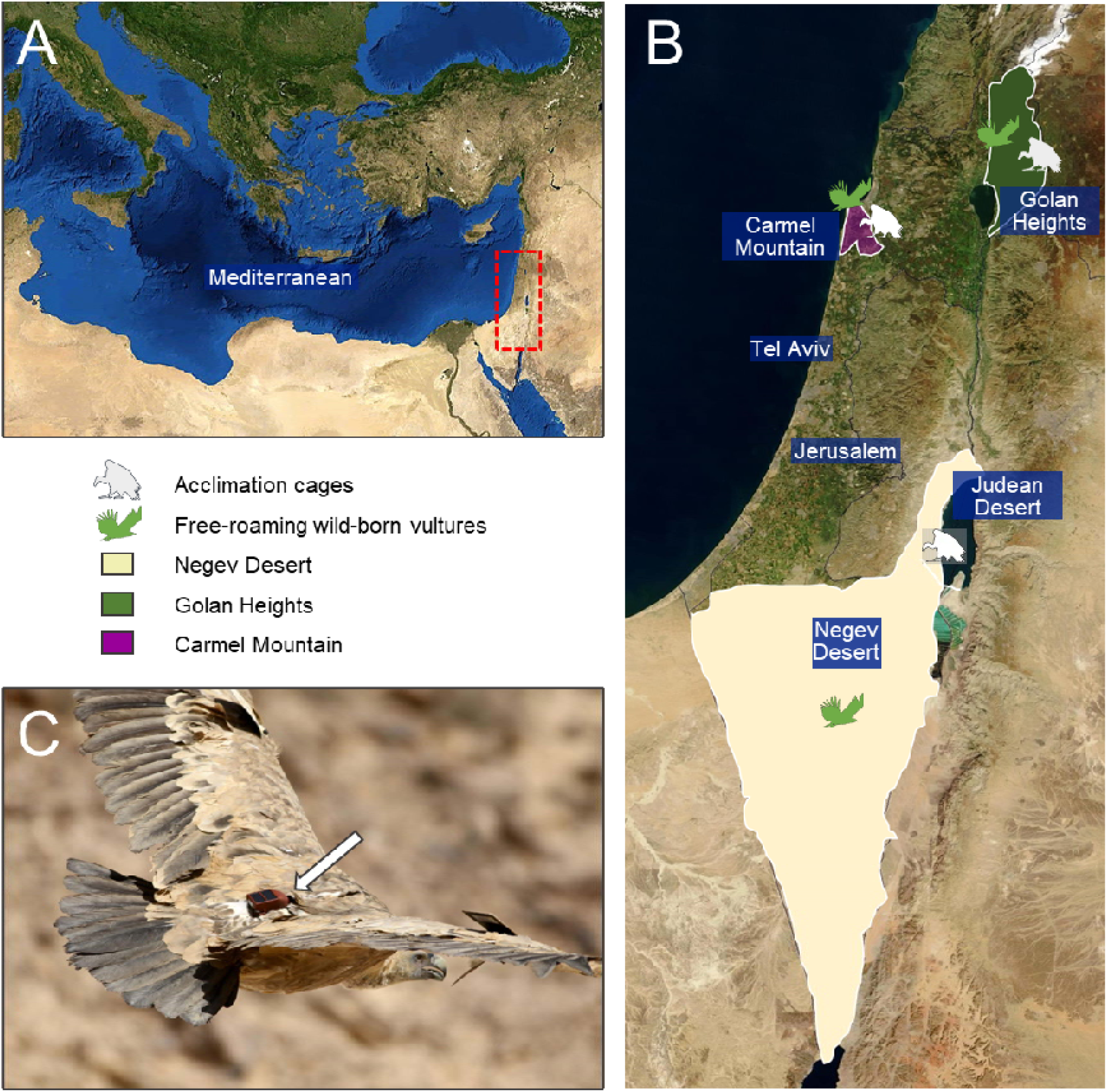
Study regions and sampling locations. (A) A regional map of the Mediterranean area in which the study region is indicated by a red rectangle. (B) Griffon activity areas within Israel: Negev and Judean deserts in yellow, Carmel mountain in purple, and Golan heights in green. White icons of standing vultures are sites of the acclimation cages, and green icons indicate that free-roaming wild vultures occur at all three regions. The green icon of a flying vulture in the Negev desert is at the site where free-roaming birds are trapped. (C) Griffon vulture with GPS transmitter indicated with a white arrow (photo credit: Tovale Solomon).

We analyze the success of the reinforcement program during the years 2017-2020, investigate possible causes for failures, and the determine the efficiency of mitigation steps. We use griffon GPS-tracking data to 1) analyze the short-term post-release survival at the three release sites in Israel; 2) examine the influence of several internal and external factors on post-release survival (e.g., weather conditions, movement behavior, pesticides, disease, and obstacles to flight that may lead to collisions); 3) evaluate whether changes in release protocols implemented following insights from 1&2 (2020 onwards) improved griffons’ short-term post-release survival. Considering the pronounced differences in climate between the northern and southern release locations in Israel (Mediterranean at the Carmel and Golan Heights sites vs. arid at Judean desert), and the extreme weather conditions experienced by griffons in the latter, we predicted that external factors, especially weather, will be a major cause of mortality in the south of the country. Furthermore, since released griffons are of Mediterranean origin, either imported from other countries (Mostly Catalonia, Spain) or captive bred in the northern parts of Israel, we predicted that those released in the desert would show higher sensitivity to the local conditions. Finally, we predicted that this mismatch between the behavior (or local adaptation) of released individuals and the harsh environmental conditions will lead to particularly high physiological stress and mortality at the desert site and that this negative effect can be mitigated by changing the release protocol to release vultures in cooler conditions. Below we describe the methodology of the translocation program and how changing wildlife management protocols facilitated a reduction in early mortality at the desert release site.

## 2. METHODS

### 2.1. Captive breeding, import, and release protocol

To overcome the rapid decline of the Israeli Griffon vulture population, captive breeding of griffons has been active in Israel since 1989. Captive-bred griffons are offspring of individuals who were brought into captivity after sustaining an injury in the wild and could not be released. Eggs are artificially incubated at the Jerusalem Zoo, to encourage the laying of a replacement clutch and maximize hatching success by offering ideal incubation conditions. The chicks are then reared either by griffon pairs at the breeding facilities (either their biological parents or a fostering pair) or using hand rearing for the first 4 months. After this period, the chicks are moved to a large aviary with other griffons at the Hai-bar Carmel breeding facility which is situated in a Mediterranean climate (32.75 N, 35.01 E). In total, 154 captive-bred griffons have been released between 1993-2023, with the first breeding for the released griffons sited in 1999. To increase population size and diversify its genetics, griffons were imported from Cyprus (n=3), Armenia (n=9), and Spain (n=115) between 2015 and 2019. All translocated griffons from other countries were quarantined at the Jerusalem Zoo for over one month to prevent the transmission of disease. Of the 115 Spanish vultures, 74 wild-born griffons, were rehabilitated in the Vallcalent rehabilitation center in Catalonia, Spain, and were released in Israel between 2015 and 2022. The remaining imported Spanish griffons, that were not released due to age and physical condition were integrated into the Israeli captive breeding program. For acclimatization, sub-adult griffons (1-4 year-old) from both the breeding program and imported griffons were moved to large aviaries at their release location 1-24 months before release, and all were released in a “soft-release” method, opening the cage and allowing them to leave and return to the adjacent feeding station. All release sites are less than 400 meters from a supplementary feeding station.

### 2.2. Study sites

The griffons in this study were released at three sites: 1) Carmel mountain (n=17, 32.75 N, 35.01 E), 2) Golan heights (n=31, 32.9N, 35.73E), 3) Judean desert (n=28, 31.4N, 35.4E) (**Fig. 1**).

The climate in the Carmel mountain and Golan Heights is sub-humid, mild Mediterranean climate with a mean annual rainfall of 400-1000mm and a hot and dry summer (Kutiel & Naveh 1987; Wittenberg et al. 2007), with average maximum summer temperatures (± SD) of 30.08°C ± 1.67 and 38.01 ± 2.41 (in August in Haifa University and Gamla Meteorological station, respectively). The Judean desert is a rain-shadow desert, located in the south-east of Israel. It is extremely arid, with a mean annual precipitation of approximately 90mm, and maximum summer temperatures averaging 40.7°C ± 2.62 (in August in Ein Gedi Meteorological station 2015-2022, Israel Meteorological Service 2023). The Negev desert is connected to the Judean desert and is where most of the free-roaming griffons in Israel reside. It is arid, with mean annual precipitation between 200mm in the north and as low as 24mm in the south and average maximum summer temperatures of 37.23°C ± 1.71 (in August in Sde-Boker Meteorological station, 2015-2022).

### 2.3. Morbidity and mortality data

We compared the major causes of morbidity and mortality among ‘wild-born’, captive-bred (hereafter ‘breeding program’), or ‘imported’ griffons, released at different sites between 2016 and 2020. We obtained data on injured and dead griffons from the Wildlife Hospital of Israel (https://www.wildlife-hospital.org.il/) and the Avian Pathology Department of the Kimron Veterinary Institute considering injured vultures as mortality events because untreated griffons would have not survived independently. Our dataset for morbidity and mortality includes free-roaming griffons found alive or recently dead, including griffons without GPS tags: 49 wild-born griffons, 38 griffons released in the north of Israel at the Carmel mountain and Golan Heights, and 24 released in the Judean desert. The griffons released at the Carmel and Golan were analyzed together as they were found at similar locations, and had similar fates. We used six categories for cause of death: (1) ‘Poisoning’ - including pesticides (mostly Choline-esterase inhibitors; see Anglister et al 2023), lead and veterinary drugs; (2) ‘Persecution’ – including gunshot; (3) ‘Infrastructure’ – which accounted for collisions with power lines and electrocution; (4) ‘Illness’ - griffons with signs of disease; (5) ‘Other’ – a small subset of mortalities from other identified factors such as bite wounds, dehydration, emaciation; and (6) ‘Unknown’ - cases with no confirmed cause of morbidity or mortality. To define the cause of death, several methods were employed, including x-rays to identify gunshot pellets (persecution), and fractures to indicate trauma (e.g., from infrastructure); blood or organs (liver, kidney, gut content) were analyzed for poisons (to identify poisoning, see Anglister et al., 2023); clinical/post-mortem hematology (including biochemistry and blood smears) as well as PCRs for pathogens (Newcastle disease virus, West Nile virus, and Avian influenza), and histopathology to identify the disease (illness).

### 2.4. Animal captures and GPS-tracking data

As part of the ongoing griffon monitoring by the INPA, captures of free-ranging, wild-born individuals occur in the Golan Heights, Carmel mountain, and Negev desert (30.86 N, 34.8 E). Walk-in traps are placed at active supplementary feeding sites and baited with large mammal carcasses during the non-breeding period (September-November). Approximately 100 griffons are captured yearly, including many re-captured. All griffons are tagged with wing tags and leg rings for individual identification and inspected externally for body condition and any signs of new injury or disease. Age is determined from feather molting for birds younger than 4 years old, or from known history (e.g., captive-raised or capture history). Feather samples are collected for genetic classification of sex (Karnieli-Vet Ltd., Kiryat Tiv’on, Israel). Blood, tracheal, and cloacal swabs are collected for a subset of captured individuals.

To quantify griffons’ movement and behavior, a subset of the captured wild griffons, as well as most translocated ones are tagged with GPS transmitters. We used Ornitrack 50 3G GPS-GSM transmitters (Ornitela, Lithuania). These transmitters weigh 50 grams (<1% of the griffon’s body mass), and are attached using a Teflon leg-loop harness, with a silicon thread inserted in the Teflon (**Fig. 1C**). Transmitters are programmed to collect a GPS position and tri-axial acceleration every 10 min, as well as the altitude above sea-level and ground-speed. Importantly for our prediction regarding the effect of weather conditions on vulture behavior and survival, these transmitters have an on-board temperature sensor that measures the ambient temperature that the animal is exposed to. As the tags transmit the data via the cellular (GSM) network, the griffons can be followed in near real-time, allowing immediate actions of INPA rangers when needed (e.g., rescuing griffons that exhibit signs of weakness or death). Between the years 2015- 2019, a total of 61 imported and breeding program-born griffons were released with GPS tags (Carmel mountain N=17, Golan Heights N=31, and Judean desert N=13). We compared their survival with the survival of 73 wild-born griffons of the same ages (sub-adults, 2nd-4^th^ calendric year), tracked in the Negev desert for at least 90 days after GPS fitting (hereafter, early phase).

### 2.5. Data analysis and statistical methods

All data processing and analysis were implemented in R (R Core Team 2018). The Generalized linear mixed models (GLMMs) described below were run with the ‘*glmmTMB*’ package for model analysis (Kristensen et al., 2016; Brooks et al., 2017) and their assumptions were validated using the ‘*DHARMa* ’ package for diagnostics of the residual of the models (Hartig, 2022). Competing models were compared through Akaike’s Information Criteria (Anderson et al., 2000). The code is available in Supplementary Materials.

#### 2.5.1 Griffon’s survival at different release sites

We used multivariate Cox proportional hazard regression (Muenchow, 1987; Hamilton et al., 2010) to study if vulture survival during the early phase after release was impacted by release site location (Carmel-Golan vs. Judean desert), season (summer: May-September or winter: October-April), age (juvenile, sub-adult, or adult), sex, and number of days in the acclimation cage (Fox, 2008). Tracking duration (days until last known status), and final status (dead, alive, or censored – i.e., unknown) were recorded for each individual. Some (N=20) of the griffons were rescued by INPA rangers after release if the GPS data and accelerometer data showed they were not flying (usually due to problems with acclimation after release), these were considered dead in the survival analysis and movement analysis, as they would not be able to survive in this condition in the wild and they were sent back to rehabilitation in the Breeding program facility (these are referred to as “caught” hereafter). The analysis was done using the ‘Coxph’ function in the R-package ‘*survival*’ (Therneau & Grambsch 2000). The best model for survival was plotted using the function ‘ggsurvplot’ and the hazard ratios were compared using the function ‘ggforest’ in the R-package ‘*survminer*’ (Kassambara et al., 2017).

#### 2.5.2. Analysis of factors influencing the success of translocations

We explored factors related to translocation success in the Judean desert, especially the very high mortality during the first 10 days after release, hereafter ‘initial phase’. We excluded the day of release itself from the initial phase. Because griffons can survive for more than two weeks without feeding (Bahat 1995; Prinzinger et al., 2002; Spiegel et al., 2013), examining the initial phase allowed us to exclude starvation as a cause of death and focus on the role of pesticides, collisions, pathogens, etc. We compared the cause of death (or injury), and the proportion of griffons with an unknown cause of death, between wild-born griffons and released griffons using a chi-square test.

We then examined if the high environmental temperature in the Judean desert could be connected to low survival in the initial period. We recorded the maximum ambient temperature at a local meteorological station (Ein-Gedi, Judean desert meteorological station) (Supplementary material **Table S1**) on the day of the vulture’s death (before time of death as detected by transmitters). This data was compared to the maximum temperature recorded by the GPS transmitter at the same time and day for both released (n=6) and wild-born free-roaming griffons (n=6) in the Judean or Negev deserts (one point per individual). We used a GLMM with a gamma distribution to compare the temperatures detected by the tags of released and wild-born vultures, using the date of death as a random factor.

#### 2.5.3 Analysis of Movement Behavior

Focusing on the Judean desert, where mortality was the highest, we studied the behavior of released griffons by comparing their flight performance with wild-born griffons active locally, and of similar age (imported N=14, breeding program N=7, wild N=36). A vulture was considered to be flying if at least two consecutive GPS fixes had a ground speed of over 4m/s. Specifically, we quantified the likelihood of flying each day (true if the griffon flew for 1 hour or more during a given day, false otherwise) and maximum displacement for the subset of ‘flight’ (true) days (measured as the distance between the first GPS fix of a day and the most distant point from that location in the daily trajectory). For both analyses, we only included griffons with a minimum of 2 days of GPS-tracking, with at least 30 GPS points per day. To ensure that we avoid bias due to the unique behavior of long range forays of griffon (Spiegel et al., 2015) we included only griffons that were tracked within Israel or the vicinity (29.55< latitude< 31.76, 34.29<longitude<35.60), leaving us with eight released griffons to study. We aimed to compare the flight performance of released griffons and wild-born free-ranging griffons of the same age, during the same dates, and within the same geographical area. However, because only three wild-born individuals from the Judean desert with active GPS tags could be matched for time and locations with the released individuals we used GPS data from wild-born griffons tracked in other years (2020-2023; N = 33) at the same dates but different years and similar locations as the released individuals from 2015-2019. To calculate movement behaviors, we used the R packages ‘*geosphere*’ (v1.5-18; Hijmans, 2022), ‘*raster*’ (v3.6-23, Hijmans, 2023), ‘*move*’ (v.4.2.4, Kranstauber, et al., 2023), and ‘*mapproj*’ (v1.2.11, McIlroy 2023). We analyzed effects on movement with a GLMM with a binomial distribution (link-logit function) with flight/no-flight days (true/false) as the response variable, and “released vs wild-born”, season, origin (“imported”, “breeding program”, or “wild”), and their interactions, as predictors, as well as individual identity and date of release as random effects. Similarly, a GLMM with a gamma distribution and the same predictors was used to examine the effects on maximum daily displacement. Due to a low sample size of captive-bred griffons that did fly within the initial phase (first 10 days after release), ‘origin’ was not included.

#### 2.5.4 Evaluation of implemented changes in the release protocol

In light of the high griffon mortality rates soon after release between 2017 and 2019, and the preliminary analyses suggesting high temperatures as a key factor enhancing deaths (see result), in 2020 all Judean desert griffons were released in the winter rather than in the late spring or summer. To assess if this change in protocol influenced the survival of the released individuals, we used multivariate Cox proportional hazard regression to compare the survival of griffons released in 2017-2019 (N=9 released in summer, N=4 released in winter), to the survival of griffons released in 2020 (N=15, all in winter).

### 2.6. Ethical aspects

The study was approved by the Israel Nature and Parks Authority (permit #42166). All samples were collected as part of annual health inspection, tagging, and telemetry tracking for monitoring and management, and no special designated captures were made for this study

## 3. RESULTS

We tracked 61 released griffons for 49±34.8 days (Mean±SD; range 1-90), and a total of 2967 days. These griffons include 17 individuals released at the Carmel mountain (63.6±32.7; total: 954 griffon-days, April 2016-February 2019); 31 griffons at the Golan Heights (54.0±33.1; total: 1724 griffon-days; December 2015- July 2019), and 13 at the Judean desert (45.8±36.4; total: 289 griffon-days; December 2017-June 2019). Another 15 griffons were released in the Judean desert during the cooler months of the year October-April 2020 (55.1± 34.5; total: 1329 griffon-days). We compared released griffons’ survival to 73 wild-born griffons that were captured and tagged in the Negev desert in the south of Israel (October 2015-November 2022). The wild-born griffons were each largely tracked for the entire 90 days, totaling 6035 days.

### 3.1. Griffon’s survival at different release sites

Our first goal was to compare the short-term post-release survival of griffons at three release sites. Out of 16 models included in our comparison, we found that the best model explaining survival during the early phase in 2017-2019, with an AIC weight of 0.73, included the location and the season of release (see, Supplementary **Table S2** for model ranking and **Fig. 2A** for effects of the top-ranking model). The highest survival rate was for griffons released in the Carmel during the winter and the lowest survival rate was for griffons released in the Judean desert in the summer (**Fig. 2A**). This difference, expressed as the high Hazard ratio (Hr=6.06, P<0.001) for vultures released in the Judean desert compared to those of the Carmel, reflects the very high observed early mortality during the initial phase of griffons released in the Judean desert. Griffons released in the winter had higher survival rates compared to griffons released in the summer (Hazard ratio=0.46, P<0.05, Supplementary **Fig. S1**), a result that supports the comparison of later releases described below. We did not detect significant effects on griffon survival of sex, origin, or the number of days in the acclimation cage before release.

**Fig. 2.**
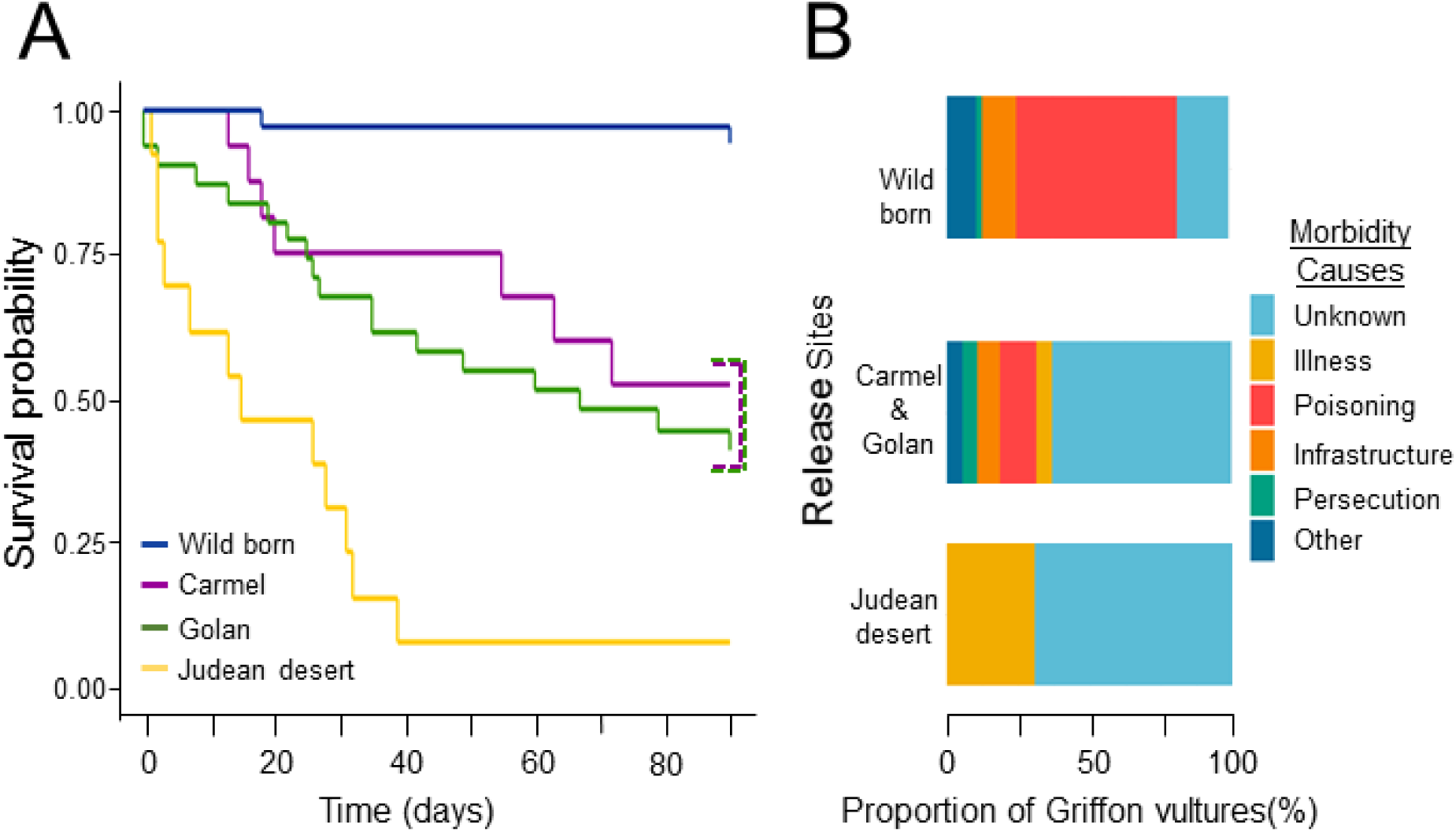
Survival and causes of mortality of released and wild griffons during the early phase (first 90 days). A. Proportion of individuals surviving over time after release in the Carmel Mountain (N=17, purple), the Golan Heights (N=31, green), the Judean desert (N=13, yellow), and wild-born griffons post-capture (Negev desert; N=75, dark blue). Mortality was significantly higher in the Judean desert compared to all other sites and mortality in the Carmel and Golan was higher than in the wild-born population, but not significantly different among these two sites**. B.** Causes of morbidity and mortality of griffons between 2016-2020. Proportion of different morbidity and mortality causes for wild-born griffons (N=49), griffons released in Northern Israel (Carmel mountains and Golan heights N=38), and griffons released in the south of Israel, in the Judean desert (N = 24).

We found that in 2017-2019, seven out of 13 griffons (54%) released at the Judean desert site died within the initial phase post-release (including one individual that was ‘rescued’ in bad condition). Similarly, within the early phase, six additional individuals from this site either died (n=1) or were rescued (n=5) in bad conditions and brought back to the breeding program facilities for rehabilitation. Ultimately, the sole survivor from the early phase died within less than a year (**Fig. 2A**). In sharp contrast, out of 73 wild-born griffons used as control (accounting for season, age, and region) for a similar 90 days’ period after GPS-tagging, none were confirmed dead, and only one was rescued within 18 days. Incomplete tracking periods reflect five GPS tags that stopped transmitting prematurely (the individuals were observed directly in some cases).

In the Carmel mountain, only one individual died during the initial phase (∼6%, suspected from a viper bite). In the early phase, five griffons died (29%) - one was rescued, and four suffered from GPS failures and could not be rescued. Two Carmel-released griffons survived the early phase but were injured during the first-year post-release, and only four individuals survived for more than a year in the wild. In the Golan heights, four died (13%) and one (3%) was rescued during the initial phase. During the initial phase, another four individuals died, seven were rescued (23%), and three GPS tags stopped transmitting (10%). Seven griffons survived the early phase but died or were rescued within the first year. Only 4 survived for more than a year.

### 3.2. Factors affecting griffons’ post-release survival

Given the exceptionally high mortality rates of griffons released in the Judean desert, our second goal was to identify the morbidity and mortality causes of this group compared to individuals at the other release sites and wild-born griffons. We found a significant difference in the proportion of unknown causes of death between wild-born griffons and released griffons (Chi-square=22.175, df=2, P<0.001). Only 18% of the wild-born griffons died of unknown causes. In contrast, for the released griffons, we could not ascertain the cause of death for 63% of the griffons released in the Carmel mountains and Golan heights and 69% of those released in the Judean desert. Of the griffons that were rescued alive, some showed signs of illness (5% in the Carmel and Golan and 31% in the Judean desert), such as weakness, tremors, and circling. However, no pathogens or toxins were found upon examination of biometric samples (**Fig.2B**).

After excluding the above-mentioned causes of death, we investigated how movement behavior and environmental conditions contributed to post-release mortality in the Judean desert by examining the GPS-tracking data. We found that the temperature measured by the GPS transmitter of released griffons that died in the initial phase was significantly higher (average 51°C) than the maximum ambient temperatures recorded at the weather station (average 38°C) or the temperatures recorded on wild-born griffons GPS tags (40 °C) (Wald test, Z=-6.9, P<0.001 **Fig. 3A**). Given the spatial and temporal and spatial proximity of wild-born and released vultures, this difference cannot represent variation in broad climatic conditions. Instead, they likely reflect differences between griffons in their ability to use behavioral thermoregulation, for example, by flying to cooler/well-vented locations or identifying shaded microhabitat.

**Fig. 3.**
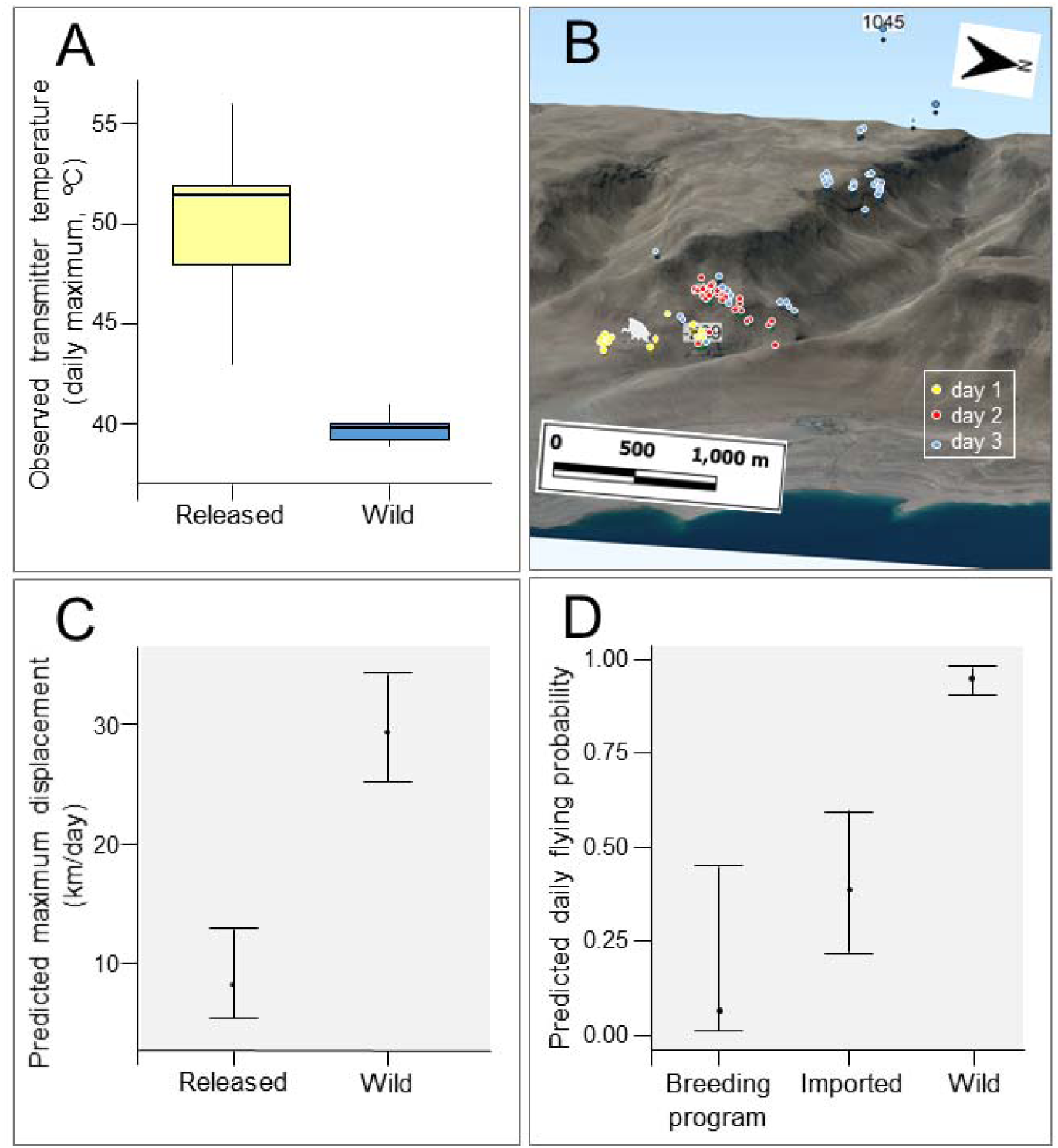
Comparing ambient temperature and movement behavior between released and wild-born griffons. **A.** Maximum daily temperature experienced by GPS-tracked released (N=6, yellow) and wild griffons (N=6, blue), as measured by the transmitter. Boxes extend from the 1st to 3rd quartile, the line reflects the median and the whiskers are 1.5 X the interquartile range **B.** Spatial positions of one imported released griffon during the first three days post-release (April 13^th^-16^th^, 2018). Each color represents a different day and points are overlaid with a 3D representation of the release site topography. In this example the griffon spent the first two days near the release aviary (griffon icon), only managing to go to higher-ground on the third day. **C**. Maximum daily displacement for days on which griffons flew, for released (imported and captive-bred grouped) and wild-born griffons. **D.** Probability of flight on a given day, by griffon origin: captive-born, imported, and wild-born. Error bars here and in panel C indicate the 95% confidence interval.

In agreement, we found that wild-born and released individuals differed in their movement behavior during the initial phase. The best fit (AICc weight of 0.38) model explaining the probability to fly included the origin of the griffons, with significant differences between wild-born and both imported (Wald test: t=-7.314, P<0.001) and captive-born griffons (Wald test: t=- 4.182, P<0.001) (Supplementary **Table S2**). The proportion of ‘flight’ days was 0.059±0.114 (estimate ±SE) for captive-born griffons, 0.381 ± 0.097 for imported (rehabilitated) griffons, and 0.951 ± 0.02 for wild-born griffons. These differences suggest that many released griffons rarely flew after release, especially those that are originally captive-born (i.e., with no prior experience in flight; **Fig. 3D, 3B).** Similarly, maximum displacement (excluding ‘no-flight’ days) was significantly shorter for released griffons that flew (8.44±1.84 km) than for wild-born griffons (29.63±2.36 km) (**Fig. 3C**, Supplementary **Table S3**). This finding supports the notion that the poor flight capacity of released griffons contributed to their inability to behaviorally avoid the extreme weather conditions in the summer of the Judean desert.

### 3.3. Evaluation of changes to the release protocol

From January 2020 through 2021 all griffon releases in the Judean desert were conducted during the cooler winter months (October-April) instead of during the hot summer. This change allowed us to determine if griffons’ inability to thermoregulate to the extreme heat was indeed a major cause of mortality, as suggested by the analyses above. Of 16 griffons released in winter three died (19%) during the early phase, and one was rescued (6%), two GPS tags did not transmit (12%), and nine individuals survived the first 90 days. At least six (38%) of these griffons survived for more than a year. Changing release timing from summer to winter significantly improved the survival of released griffons by 76% (Hazard ratio=0.24, P=0.005; **Fig. 4**, Supplementary **Fig. S2)**.

**Fig. 4.**
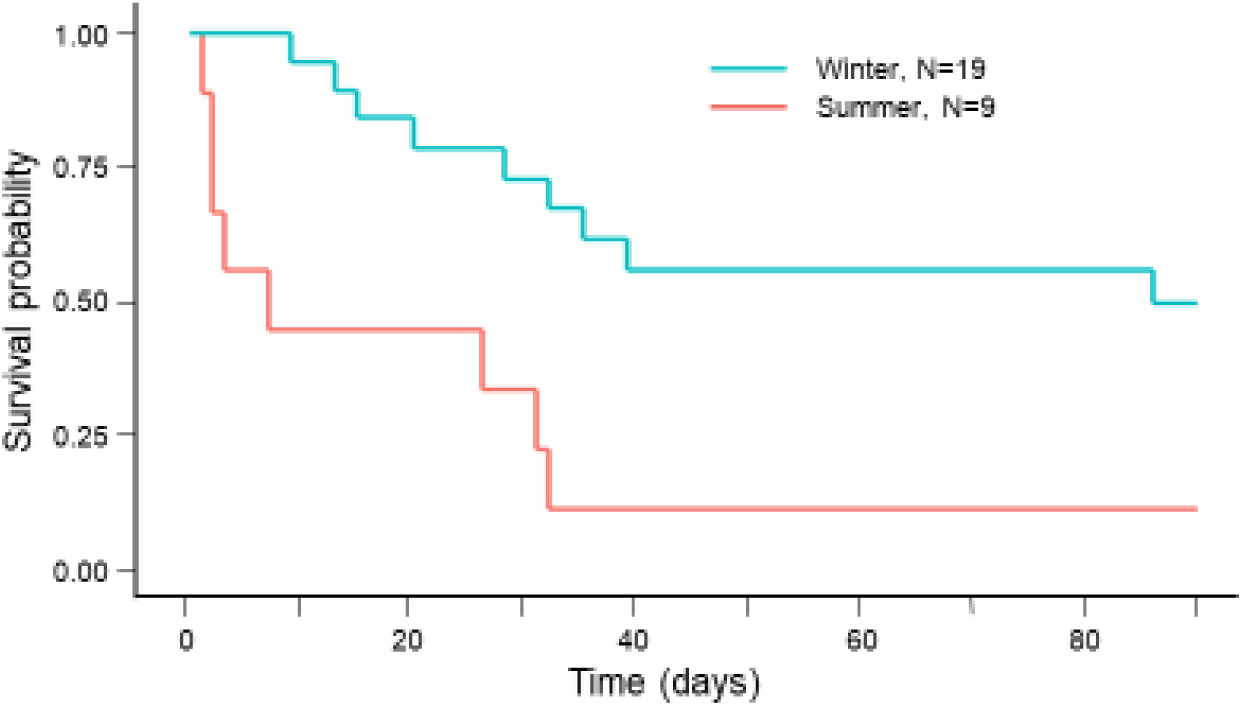
The survival rate of griffons during the early phase (90-days) post-release at the Judean desert. Between 2017 to 2019 nine griffons were released during the summer months (May-Sep, red). Following the insights from the preliminary analysis reported above, release protocols were modified to the cooler winter months (October-April, turquoise) for the 19 griffons released through 2020 and 2021

## 4. DISCUSSION

Here we studied factors affecting the survival of translocated griffon vultures and found that the survival of released griffons varied significantly across sites and seasons. Griffons released at the Judean desert site during the summer months experienced the lowest survival rate during the early phase (the first 90-days from release), with exceptionally high mortality during the initial phase (the first 10-day from release). Nevertheless, despite this high mortality rate, the specific cause(s) of death remained elusive until now. Intriguingly, the higher temperatures measured by the GPS tags of these released individuals suggest a potential link between heat exposure and low survival. Moreover, we found that released griffons (especially those that were captive-bred with no prior flight experience) had a lower probability of flying and had shorter flight distances than wild-born griffons. Together, these findings suggest that the limited flight capability of captive-bred (and imported) griffons prevented them from avoiding the extreme temperatures (e.g., by flying to shade), contributing to their elevated vulnerability in hot conditions. Free-roaming griffons and those released at other sites, and/or during the winter, were less exposed to these hot conditions and survived better. Consequently, release protocols were changed and subsequent Judean desert releases (during 2020 and 2021) were conducted during the winter. In agreement with our conclusion, survival during these later releases increased, particularly during the critical period of the initial phase. Below we discuss the factors affecting the survival of released griffons, compare them to factors identified in other systems and summarize how these insights can be translated to conservation actions.

### 4.1 The effect of release site on short-term survival

Notably, survival in Israel (across all sites) was much lower than survival reported for translocation of griffons in France, where the estimated annual survival is 0.987 for adults and 0.858 for juveniles (Sarrazin et al., 1996; Sarrazin & Legendre 2000). Further, in contrast to previous research on vulture reintroductions (Sarrazin & Legendre 2000; Germano et al., 2014; Efrat et al., 2020; Thomas et al., 2023), neither sex, age, number of days in acclimation, or origin of the griffons affected the survival during the first three months in our study. This finding suggests that other factors may impact post-release survival during the initial phase.

High mortality occurred within the initial phase with 46%, 13% and 16% of released griffons dying in the first 10 days in the Judean desert, Golan and Carmel, respectively. These results align with the critical period for survival found in translocations of mammals (Maran et al., 2009; Devineau et al., 2010; Hamilton et al., 2010). In general, reintroduced animals acclimate and acquire knowledge about their new surroundings during the first few weeks. During this critical period, they explore their new sorrounding and develop the skills needed to survive, such as flight and knowledge about resources (Eliassen et al., 2007; Pinter-Wollman et al., 2009; Maor-Cohen et al., 2021). This initial phase (and beyond) requires relying on existing energetic reserves, and therefore the animals’ initial physical condition is crucial for survival (Eliassen et al., 2007; Berger-Tal et al., 2014). Nevertheless, even with low survival expectation in mind, post-release survival in the Judean desert was alarmingly low in comparison to other release sites in our study (and to other translocation studies; e.g. Sarrazin & Legendre 2000; Gouar et al., 2008b; Efrat *et al.,* 2020). It was particularly surprising to find such differences in survival between the locations as the distance between the various sites ranges between 70-200 km, and can be easily crossed in a day by a griffon (García-Ripollés et al., 2011; Monsarrat et al., 2013; Spiegel et al., 2015). However, because we observed low dispersal of the griffons after release, we suspect that differences in the survival between the locations can be explained by spatial heterogeneities of threats to the griffons as suggested by Le Gouar et al. (2008) and Efrat et al. (2020) and by the sharp climatic gradient among sites, as we explain below.

The season of release affected post-release survival in all release locations (including the Carmel and Golan), with winter releases having 54% higher survival compared to summer releases for the early phase. These results are surprising considering that the opposite was found for one-year survival in the Carmel and Golan sites previously (Efrat et al., 2020) with higher survival for releases in the summer months. Efrat et al. (2020) suspected that in the Mediterranean climate griffons would be able to use thermals for soaring and gliding flights during the summer, and reduce energy loss compared to flapping flights in winter. The Judean desert, however, is a very arid environment with summer temperatures exceeding 40°C (Israel Meteorological Service 2023), thus the challenges in this environment are different from those in more temperate climates. Desert climates are characterized by water and food and shade under vegetation scarcity, and extreme heat during the summer (Kutiel et al., 1995; Shoshany et al., 1996). In addition, there are extreme differences in temperature between direct sun and shade (Hardman & Moro 2006), and these abiotic conditions may challenge released griffons that are naïve to their surroundings.

### 4.2 Morbidity and mortality causes of released and free ranging griffons

Beyond the spatial heterogeneity among release sites, our findings of causes of morbidity and mortality in wild-born griffons were consistent with previous research of vulture populations worldwide (Ogada *et al*. 2012; Buechley & Şekercioğlu 2016; Anglister et al., 2023). Expectedly, poisoning (mainly pesticides, but also heavy metals and anti-inflammatory drugs) along with electrocutions and collisions with infrastructure were the main risks for wild-born griffons. However, released griffons died for other reasons, especially those in the Judean desert. Although almost all of the griffons were equipped with GPS that facilitated quick recovery (Acácio et al., 2023) and despite intensive tests for pathogens and toxins, we could not establish the cause of death in most of these cases (over 60% of all released). Some of the griffons did show weakness, tremors, and circling behavior, however, no pathogens, toxins, or conclusive pathological or clinical findings were identified. In contrast to previous studies implying that translocated griffons suffer from the same risk factors as the local population (Hamilton et al., 2010; Bubac et al., 2019), this was not the case for individuals in the Judean desert. Thus, we argue that while risks might be similar to the native population for released individuals that made it through the early “acclimation period”, during this early period itself the risks are substantially different, reflecting the limited physical capacity and knowledge of the released individuals. This deviation is expected to be particularly pronounced for species that rely on locomotion modes that cannot be developed in captivity, such as soaring flight for our griffons, impairing their ability to obtain resources, and avoid enemies and difficult environmental conditions.

In agreement with this explanation, starvation was shown to play a major role during the early post-release survival of several reintroduced species, either due to a lack of knowledge of where to find food or ability to obtain it (Snyder et al., 1996; Griffin et al., 2000; Yoda et al., 2004; Hamilton et al., 2010; Feenders et al., 2011; Whiteside et al., 2016). While starvation could explain deaths after two or three weeks (in the Carmel and Golan heights), it is unlikely because all release sites were close to supplementary feeding stations (<400m). Further, because griffons are known to be adapted to fasting for over two weeks (Bahat 1995; Prinzinger et al., 2002; Spiegel et al., 2013), starvation cannot explain high mortality during the initial phase at the Judean desert, supporting our explanation of exposure to extreme heat stress (hyperthermia).

### 4.3 Temperatures experienced by released vs. wild-born griffons

We hypothesize that translocated animals may be unable to cope with unfamiliar environments; in our study these are extreme heat conditions. For introduced vultures, this means being unable to soar to cool down or find shade, similar to fledglings of free-ranging vultures (Harel et al., 2016; Rotics et al., 2016; Buechley et al., 2021). For example, harsh environmental conditions, such as extreme temperatures, have a negative effect on the survival and translocation success in other birds (Altwegg et al., 2006; Nevoux et al., 2008; Tavecchia et al., 2009; Hardouin et al., 2014). The high ambient temperatures measured by the vulture-borne transmitters imply that released griffons failed to avoid the heat, resulting in direct and fatal exposure to solar radiation in the hours preceding death (hyperthermia). Because the pathophysiology of hyperthermia in birds has not been well characterized, direct identification in clinical examination is not simple, and it is difficult to identify in necropsy.

Hyperthermia may seemingly contradict griffons’ morphological adaptations for regulating heat exchange (Arad et al., 1989; Bahat 1995; Ward et al., 2008) allowing them to occupy a very wide geographical distribution and thermoneutral zone (Bahat 1995; Prinzinger et al., 2002). Nevertheless, we note that normally griffons can adjust *behaviorally* to extreme weather conditions. Bahat (1995) showed that flight alone can reduce the measured surface temperature of griffons by 4-7°C compared to griffons on the ground (measured in an ambient temperature of 30-34°C during the summer in Northern Israel). This result highlights the importance of flight as a cooling mechanism for griffons, in addition enabling the vultures to reach shaded/vented microhabitats. Unlike local, free-ranging griffons dwelling in the same season and region, those released in the summer of the Judean desert were exposed to temperatures of 43-56°C (which may exceed the thermos-neutral zone, known to be between 5-35°C; Bahat, 1995) likely leading to hyperthermia. Reduced flight capacity (both in probability of flight and in distances) of released griffons was more pronounced in the captive-bred griffons compared to the imported ones. Indeed, captive-bred vultures were raised in an aviary and lack any prior large-scale soaring flight experience, while the imported ones were wild-born (in Catalonia, Spain) and had previously experienced such flights, and were therefore less immobile.

Finally, another (system-specific) factor that might potentially contribute to the impaired flight capacity of the Judean desert-released individuals is the suboptimal topographic placement of the acclimation cage. The cliffs in this area start at ∼400 m below sea level, reaching the upper plateau at around 300m above sea level and the cage is placed at the lower part of the cliff height (276m below sea level, on the head of an alluvial fan with moderate slopes; see Fig. 3B), due to accessibility limitations. This limits the take-off ability of inexperienced vultures by offering poor uplift conditions. Indeed, rangers observed that released griffons at this site often spend their first few days climbing the cliff by foot, rather than flying (*INPA rangers*, personal observations). Therefore, it is vital to choose the conditions of the release site wisely relating to climate, season, landscape, local topography, and micro-climate.

### 4.4 The evaluation of changes to release protocol of griffons

Management protocols were updated following recommendations based on our preliminary results. From January 2020 griffons in the Judean desert were released in cooler conditions during the winter (October-April). This change to wildlife management protocols resulted in a direct increase (by 76%) in the short-term survival of griffons released in the Judean desert.

### 4.5 Study limitations

This study is not without limitations, including a relatively small sample size and a focus on short-term survival. Future research endeavors should expand on these limitations by incorporating larger sample sizes, longer study periods, more release sites (with account of elevation) and more comprehensive physiological assessments to further enhance our understanding of translocation dynamics. While these are not available in the Israeli griffon management program, they can be obtained in a comparative study with other parallel programs (e.g., in Crete, Cyprus and more). Nevertheless, this data has already helped reform conservation management decisions regarding optimal release conditions for the Griffon vultures in Israel and can be useful in translocation projects involving vultures and other species elsewhere.

### 4.6 Conservation implications

Our findings that vultures could not cope with the environmental conditions due to a limited ability to behaviorally seek safe shade spots is likely transferable to many other systems where the limited physical fitness and knowledge of released individuals can interact with early conditions in other climates (such as limited fat reserves in extremely cold climate, or finding food and water sources).

By emphasizing the importance of release site selection, season, and flight ability, our study provides valuable lessons for improving the survival and success of translocated animals. Our results exemplify that the desert climate has a different set of challenges compared to temperate regions and requires adjusting wildlife management techniques. In a world facing climate change with temperatures becoming more extreme and unpredictable, these challenges will probably become more widespread. We also highlight the need for adaptive management strategies, sharing information on failures, and incorporating multidisciplinary data analysis to ensure the long-term viability of endangered species in their native habitats.

## Acknowledgements

The conservation translocation project for Griffon Vultures in Israel is led by the Israel Nature and Parks Authority (INPA) and financed by the Porsim Kanaf partnership with support and cooperation from Keren Hayesod, Segre’ Foundation, the I. Meier Segals Garden for Zoological Research at the Tel-Aviv University, the Zoological Center Tel-Aviv—Ramat-Gan, Ramat Hanadiv Gardens, especially Y. Levy- Paz and E. Zisso and Arkia. Chicks’ rearing was done in partnership with the Jerusalem Biblical Zoo, led by N. Avni-Magen. Most of the work preparing the vultures for translocation is performed by the Hai-bar Carmel Nature Reserve team, led by A. Baron and S. Simchi, the Gamla Nature Reserve team, especially A. Maatuf, and the Ein Gedi Nature Reserve team, especially D. Shilo & L. Lev. We want to thank the Wildlife Hospital of Israel team: especially R. Elias-Ofri, for their diagnosis and dedicated treatment of hospitalized Griffons. We want to thank T. Solomon, A. Uzan, S. Cain, and many others from the Tel-Aviv University Movement Ecology Laboratory for their assistance in the field and for their valuable advice. We are indebted to A. Lublin, A. Berkowitz, I. Farnushi, S.Mashani, O. Cunaeh and M. Britzi from the Division of Avian Diseases, Kimron Veterinary Institute, for the pathological and toxicological examination of the Griffons, and to R. Lapid, R. Efrat, A. Gancz, G. Kahila-Bargal, and Z. Aisenberg for their advice. The work was funded by the NSF IOS division 2015662/BSF 2019822 (to OS and NPW). NA was supported by a stipend from Yad-Hanadiv.

## Author contributions

**NA**- Conceptualization, Methodology, Software, Formal Analysis, Investigation, Data curation, Writing-Original draft, Visualization, Validation & Supervision.

**MA**- Formal Analysis, Validation, Visualization & Writing-Review, and Editing.

**GV**- Formal Analysis, Validation & Writing-Review and Editing.

**EA**- Software, Formal Analysis & Validation

**MB**- Validation & Visualization

**OH**- Conceptualization, Resources, Project administration & Funding acquisition.

**YM-** Methodology, Project Administration & Data Curation

**RK-** Methodology, Project administration & Funding acquisition

**NPW-** Writing-review & editing, Visualization, Supervision & Funding acquisition.

**OS**- Conceptualization, Methodology, Formal Analysis, Software, Writing-review & editing, Visualization, Validation, Supervision & Funding acquisition.

