## Supplementary tables for "Extreme temperatures impede the release success of captive-bred avian scavengers"

**Supplementary Material**

I. **Supplementary Methods**

**Table S1. Table of vultures GPS tag maximum temperatures before the time of death of released griffons** according to accelerometer data compared to temperatures of GPS transmitters of paired wild-born griffons at the same time. Ambient temperature is taken from the maximum temperature per day according to the En Gedi meteorological station database (Israel Meterological Service 2023).

| Griffon I.D. | Date | Origin | Group release | Israel time | Max tag temperature | Max Ambient temperature |
| --- | --- | --- | --- | --- | --- | --- |
| T24 white | 25 July 18 | Released | 1 | 17:30 | 43 | 43 |
| Y21 white | 25 July 18 | Wild | 1 | 17:30 | 40 | 43 |
| T81 white | 13 August 18 | Released | 2 | 14:05 | 47 | 39.8 |
| K39 white | 13 August 18 | Wild | 2 | 14:05 | 38 | 39.8 |
| T80 white | 17 May 19 | Released | 3 | 14:14 | 52 | 40.2 |
| T23 white | 17 May 19 | Wild | 3 | 14:14 | 40 | 40.2 |
| T29 white | 21 May 19 | Released | 4 | 14:32 | 56 | 38.1 |
| T41 black | 21 May 19 | Wild | 4 | 14:32 | 40 | 38.1 |
| T28 white | 5 June 19 | Released | 5 | 18:18 | 51 | 41.3 |
| T22 black | 5 June 19 | Wild | 5 | 18:18 | 41 | 41.3 |
| A99 white | 6 June 19 | Released | 6 | 15:27 | 52 | 40.9 |
| K29 white | 6 June 19 | Wild | 6 | 15:27 | 39 | 40.9 |

II. **Supplementary Results**

| **Table S2.** **Factors affecting** **survival of released griffons in 2015 to 2019.** Model rankings based on Akaike Information Criterion (AICc) for predictors of 90-day survival after release. Only taking into account models with ΔAIC <4. | | | | | |
| --- | --- | --- | --- | --- | --- |
|  | K | AICc | Delta_AICc | AICcWt | Cum.Wt |
| Region and Season | 3 | 261.31 | 0.00 | 0.77 | 0.73 |
| Region | 2 | 264.26 | 2.96 | 0.22 | 0.89 |

**Figure S1.** **Hazard ratios of** **survival of released griffons.** The best model for survival using Cox proportional regression model during the first 90 days between 2015-2019 included the location and the season. Higher hazard ratios indicate lower survival probability.

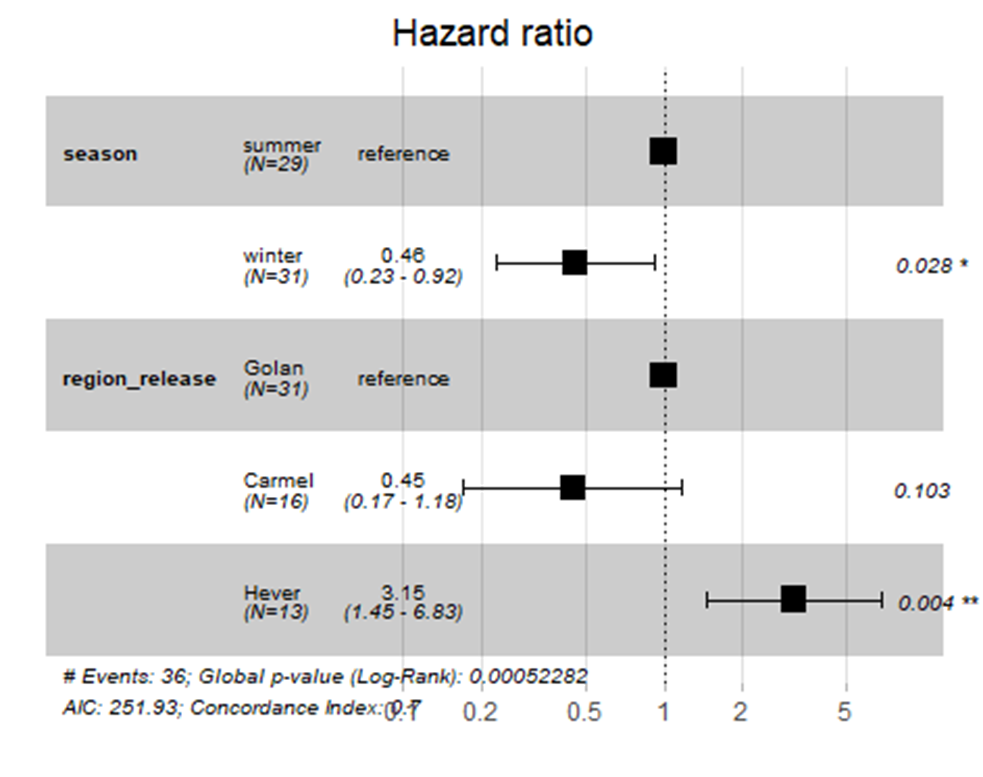

**Table S3.** **Factors explaining the probability of flying for the first 10 days post-release** for logistic regression models with individual and date of release as random factors. Model rankings based on Akaike Information Criterion (AICc) Only taking into account models with ΔAIC <4.

|  | K | AICc | Delta_AICc | AICcWt | Cum.Wt |
| --- | --- | --- | --- | --- | --- |
| Origin | 5 | 269.12 | 0.00 | 0.38 | 0.38 |
| Released *vs.* Wild | 4 | 270.73 | 1.61 | 0.17 | 0.55 |
| Origin and Season | 6 | 270.93 | 1.81 | 0.15 | 0.70 |
| Origin and Season & interaction | 6 | 270.93 | 1.81 | 0.15 | 0.85 |
| Released *vs.* Wild & Season | 5 | 272.39 | 3.27 | 0.07 | 0.93 |
| Released *vs.* Wild & Season & interaction | 5 | 272.39 | 3.27 | 0.07 | 1.00 |

**Table S4.** **Parameter estimates for the best model used for flying probability for griffons for the 10 days after release** was for origin, random effects included the individual id and date of release.

| Parameter | Estimate | SE | z | p |
| --- | --- | --- | --- | --- |
| Breeding program | -2.76 | 1.31 | -2.11 | 0.035* |
| Catalonian | 2.27 | 1.36 | 1.68 | 0.094 |
| Wild | 5.71 | 1.37 | 4.18 | 2.89e^-5^*** |

*P<0.05, P**<0.01 and P**<0.001

**Table S5.** **Parameter estimates for the model comparing released vs. wild born griffons’ maximum displacement per days** for the first 10 days after release random effects included the individual I.D. and date of release.

| Parameter | Estimate | SE | z | p |
| --- | --- | --- | --- | --- |
| Released | 2.23 | 0.22 | 10.23 | 2 e^-16^*** |
| Wild | 1.13 | 0.22 | 5.26 | 1.42e^-7^*** |

*P<0.05, P**<0.01 and P**<0.001

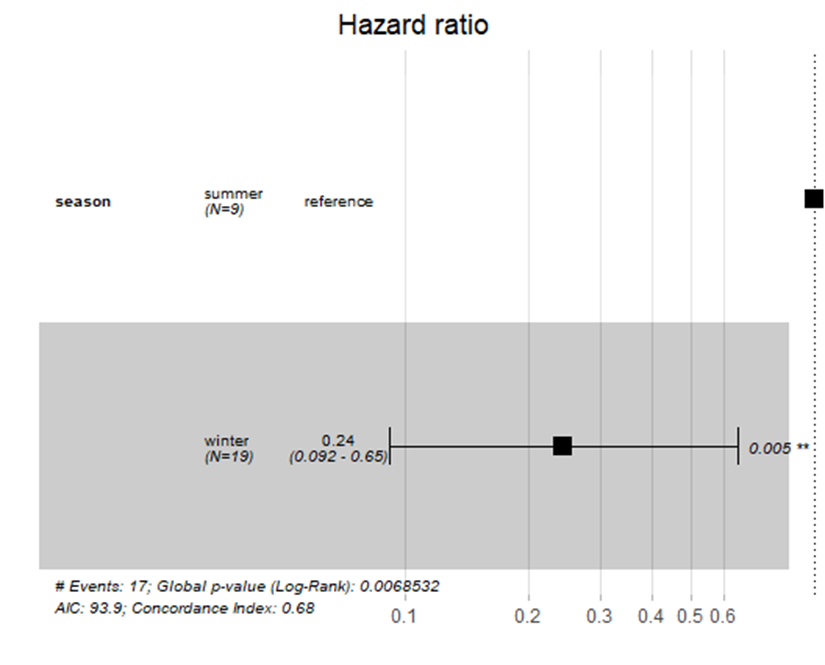

**Figure S2.** **Hazard ratios of** **survival of released griffons in the Judean desert comparing release in cooler vs. warmer months.** Higher hazard ratios indicate lower survival probability. In 2020 all releases were during the cooler months.
